# Isolation, identification and functional characterization of cultivable bacteria from Arabian Sea and Bay of Bengal water samples reveals high diversity

**DOI:** 10.1101/2020.07.31.229039

**Authors:** Shriram N. Rajpathak, Yugandhara M. Patil, Roumik Banerjee, Asmita M. Khedkar, Pawan G. Mishra, Mandar Paingankar, Deepti D. Deobagkar

## Abstract

The oxygen minimum zone of the Arabian Sea (AS) and Bay of Bengal (BOB) is rich in organic matter and is an unusual niche. Bacteria present in the oceanic water play an important role in ecology since they are responsible for decomposing, mineralizing of organic matter and in elemental cycling like nitrogen, sulfur, phosphate. This study focuses on culturing bacteria from oxygen minimum zones (OMZ) and non-OMZ regions and their phylogenetic as well as the functional characterization. Genotypic characterization of the isolates using amplified rDNA based 16SrRNA sequencing grouped them into various phylogenetic groups such as alpha-proteobacteria, gamma-proteobacteria and unaffiliated bacteria. The cultivable bacterial assemblages encountered belonged to the genus *Halomonas*, *Marinobacter*, *Idiomarina*, *Pshyctobacter* and *Pseudoalteromonas.* Among the enzymatic activities, carbohydrate utilization activity was most predominant (100%) and microorganisms possessed amylase, cellulase, xylanase and chitinase. A large proportion of these bacteria (60%) were observed to be hydrocarbon consuming and many were resistant to ampicillin, chloramphenicol, kanamycin and streptomycin. The high diversity and high percentage of extracellular hydrolytic enzyme activities along with hydrocarbon degradation activity of the culturable bacteria reflects their important ecological role in oceanic biogeochemical cycling. Further assessment confirmed the presence of nitrogen reduction capability in these cultivable bacteria which highlights their importance in oceanic geochemical cycling.

## 1. Introduction

Marine microorganisms are exposed to diverse environmental parameters including temperature, pressure, salinity and oxygen concentration [1]. Bioactive compounds discovered from marine microbes are important for biotechnological applications and have medicinal importance [2–5]. More than 15,000 chemical substances including 4196 bioactive marine natural products have been isolated from marine organisms [6] highlighting the potential and importance of cultivable marine microorganisms.

Oxygen plays an important role in shaping the aquatic ecosystem which controls the spatio-temporal distribution of all living organisms from microbes to higher organisms. Low oxygen (<20 μM) [7] regions can be observed in various aquatic ecosystems including stagnant freshwater, saline lakes [8], and marine basins with restricted circulation [8]. In marine regions of high productivity with slow ventilation rates [8], dissolved oxygen concentrations at lower depths drop to very low levels, these regions are referred to as oxygen minimum zones (OMZ).The differential solubility of oxygen and consumption via microbial-mediated oxidative processes are also known to affect the oxygen concentration in marine water.

In oceanic OMZ areas, anaerobic respiration by microbes includes use of alternative electron acceptors such as nitrate and sulfate [7]. Thus, presence of OMZ areas influences the chemical composition of the water and induces stress on organisms including alterations in the food web structure [8, 9]. For instance, total reduction of nitrogen and release of nitrogen gas by microbes removes the fixed nitrogen from marine ecosystem [8]. Similarly, sulfur (S) reduction produces hydrogen sulfide and increasing the toxicity of surrounding environment [10, 11]. The Indian Ocean, the Arabian Sea and Bay of Bengal covers the 59% of the total global OMZ area [12]. Some previous studies on OMZ have concentrated primarily on the diversity of microorganisms in waters/sediments from Arabian sea [13–15] and Bay of Bengal [16–21]. In addition, the identified cultivable bacteria during algal blooms in AS region were shown to have denitrifying capability, indicating relationship between water column characteristic and microbial diversity [22]. Additionally presence of active nitrogen cycle [23–25], sulfate reduction [26] and methane metabolism [27] from Arabian sea have been reported. In a recent study [28] where abundance of microbial genes associated with N_2_ production have been quantified which supported the presence of denitrifiers and anammox microbial population in Bay of Bengal OMZ area. BOB –OMZ is weaker due to less detritus along the western boundary [29] than AS-OMZ. Unlike Arabian sea, Bay of Bengal receives immense fresh water runoff as well as average sediment load (1.1 × 109 tonnes) [30, 31] from major rivers. Thus, overall AS-OMZ and BOB-OMZ differ in various aspects including salinity, DO concentrations and sediment load.

Denitrification is a dissimilatory process which involves the oxidized nitrogen compounds as alternative electron acceptors for energy production during very low oxygen concentration. Nitrogen oxides are reduced to gaseous products (NO, N_2_O, and N_2_) which are released, leading to a loss of fixed nitrogen to the environment. In marine coastal sediments, denitrification removes 40 to 50% of external inputs of dissolved inorganic matter[32] resulting in an unbalanced nitrogen budget in the ocean [33, 34]. Accumulation of NO and N_2_O contributes to global warming and the destruction of the ozone layer. Also the mangrove studies reveals that the anthropogenic activities lead to the bioaccumulation of different metals which in turn affects its associated biota [35]. Thus, there is a need for understanding the community of denitrifying bacteria as they influence the environmental conditions.

In addition to 16S rDNA, several functional genes were shown to be useful in the investigation of microbial communities [36–38]. Functional genes provide a resolution below species level and this approach may indicate functional diversity in the environment. An approach involving 16S rDNA alone does not appear to be suitable to investigate communities of denitrifying bacteria, as denitrification is widespread among phylogenetically unrelated groups [39]. Present study investigates the cultivable bacterial diversity from OMZ and non-OMZ areas of AS and BOB. Microorganisms have been cultured and characterized based on 16SrRNA sequencing. Further functional characterization includes the substrate utilizing properties. Hence this study deals with comparative taxonomic as well as different substrate utilization properties of cultivable bacteria form Indian OMZ and Non-OMZ areas. .

## 2. Materials and Methods

### 2.1. Sample collection and isolation of microorganisms

Water samples from OMZ and Non-OMZ regions of Arabian Sea and Bay of Bengal were collected in Apr/May 2015 (Cruise No. 340,) and in December 2015 (Cruise No. 346) Sagar Sampada, CMLRE, MOES respectively. CTD profiler was used for collection of Water samples and measuring the DO concentration.

Water samples (100μl) were spread on Zobell’s media containing 5% sea salt and incubated at 12°C in dark for 7 days. Morphologically different colonies were isolated and purified. These pure single isolates were then used for subsequent analysis including Gram staining and substrate utilization assays. Isolates were also examined for their growth at 12°C and 37°C temperature so as to find out the optimum temperature required for their growth.

### 2.2. Identification of microorganisms

In total, 40 single colonies from Arabian Sea and 31 distinct single colonies from Bay of Bengal were isolated. From all the 71 colonies genomic DNA was isolated using QIAprep® kit according to the manufactures protocol. This was followed by 16S rRNA PCR using 27F in combination with1492R/1390R/519R set of primers (Online resource 1). Total 25 μl of total PCR reaction with 1 μl DNA (50–100 ng), 1 μl each of primers (10 pmol μl−1), 2.5 μl 10× Taq polymerase buffer (Promega), 0.5 U Taq DNA polymerase (Promega) and 200 μM of each dNTPs (Promega) was set up. The PCR conditions were, 5min at 95°C for initial denaturation, 35 cycles of 30 s at 95°C, 45 s at 48°C or at 55°C, 120 s at 72°C and a final extension for 10 min at 72°C. PCR product was checked using 1% agarose gel electrophoresis and purified using Promega (A9282) kit according to manufacturer’s instructions. Sanger dideoxy method was used for sequencing all the PCR products of 16S rDNAand the taxonomic identification of each bacteria was carried out using NCBI BLAST tool.

### 2.3. Screening for ability to use different Substrates

#### 2.3.1. Enzyme Assays

All the 71 isolates were screened for extracellular production of amylase, xylanase, agarase, cellulase and chitinase enzymes. For agarase and amylase, 1% agar and 1% starch plates were used, respectively and clear zones produced by the isolates were identified using Gram’s iodine with potassium iodide staining. For cellulose, 1% CMC and for xylanase, 1% Birchwood xylan plates were used and clear zones were observed after 2/3 days using Congored. For Chitinase assay, 1% colloidal chitin plates were used and clear zones were directly observed after 3 days of incubation. Bacterial isolates which showed clear zones in plate assays were used for colorimetric quantitative assays for amylase, xylanase and cellulase. Initially, the cultures were grown in minimal medium using 1% soluble starch, 1% birch-wood xylan and 1% CMC as a sole carbon source for amylase, xylanase and cellulase, respectively. After 72 hrs of incubation, cell free supernatant was used for colorimetric assay. For cellulase, xylanase and amylase assay 20μl of 1% CMC, 1% xylan and 1% starch was added in 96 well plate with 20μl cell supernatant from each isolate. Plates were incubated for 1hr. Further 160μl of DNS solution was added in each well and incubated at 100°C for 20 min followed by reading at 570nm in multimode plate reader (Perkin Elmer). All the quantitative assays were done in triplicates and the activity was estimated using the standard graph of glucose.

#### 2.3.2. Hydrocarbon degradation

For Hydrocarbon degradation assay, isolates were grown in Bushnell-Hass medium (Himedia-M350) with kerosene as a sole carbon source. Microbes which were able to grow in this medium were considered as positive for kerosene degradation.

#### 2.3.3. Biochemical Test for Carbohydrate utilization

Carbohydrate utilization kit (HiCarbo™KB009) containing 35 different carbohydrates was used to study the utilization of different carbohydrates. The kit contains 35 wells of different carbohydrates with a negative control. 50μl of freshly grown culture was inoculated in each well and color change was observed according to the manufactures instructions.

#### 2.3.4. Sulfur Reduction Test

Barr’s medium was used to test the sulfur reducing capacity of different isolates. The medium contains different sulfate salts which turn black when utilized by any organisms. Isolates were inoculated in 10 ml of medium and kept in dark. Color change from white to black indicated the positive reaction and confirmed the sulfur reduction. *Desulfovibrio desulfuricans* was used as a positive control.

#### 2.3.5. Nitrate reduction test

Nitrate reducing ability was assessed by Nitrate/Nitrite colorimetric assay kit (780001) from “Cayman Chemicals” with slight modification in manufactures protocol. Bacterial culture was suspended in 500μl of 1X PBS buffer, lysozyme (400μg/ml) was added in each tube followed by sonication. Samples were incubated at 37°C for 1hr and centrifuged at 14,000 rpm for 20 min at 4°C. Supernatant was removed and protein concentration was quantified. Total cell protein (50 μg) for each sample was used along with 80μl of assay buffer (from the kit). Then 20μl of Nitrate standard (From Kit, concentration) was added in each well. Assay plate was incubated at RT in dark for 1hr, 50μl of Griess reagent I followed by 50μl of Griess reagent II was added in each well and colour development was observed. If pink color appeared, it indicated the presence of nitrite and the test was considered positive. In case no colour developed, pinch of zinc dust was added to the well. Development of colour after addition of zinc showed that nitrate was not reduced and the test is negative but if no colour appeared, it indicated the total reduction of nitrate and confirmed that nitrogen is released from the sample indicating positive test.

### 2.4. Antibiotic resistance

All seventy-one isolates were screened for Ampicillin, Chloramphenicol, Streptomycin and Kanamycin resistance by disc diffusion method. Bacterial culture (100μL) was spread on Zobell’s agar plate and discs were placed on the plate. On discs, 10 μL of each antibiotic (100μg/mL concentration) was added. Presence of clear zone around disc after overnight incubation indicated sensitivity to antibiotic.

## 3. Results and Discussion

### 3.1. Cultivable microbes from AS and BOB OMZ-Non-OMZ area

Initial screening of samples led to the identification of 71 distinct types of microbes (Online resource 2). Detailed sampling information of the water samples collected from AS and BOB has been mentioned in Table I. Isolates were examined for their growth at two different temperature i.e. 12°C and 37°C and it was observed that all the isolates were able to grow on both the temperature. Interestingly it was observed that at 12°C the growth was much slower than at 37°C. In future studies however, it will be interesting to understand the metabolic difference in bacteria grown at these temperatures.

**Table I.** Details of sampling site and Physico-chemical parameters of samples collected from A) Arabian Sea and B) Bay of Bengal waters.

| Location | Latitude | Longitude | Depth (m) | Oxygen concentration (ml/L) |
| --- | --- | --- | --- | --- |
| Calicut | 11.377 | 74.712 | 1000 | 0.5559 |
| Calicut | 11.621 | 74.903 | 189 | 0.28106 |
| Calicut | 11.37 | 75.128 | 96 | 0.51704 |
| Goa | 15.755 | 72.819 | 1001 | 0.3527 |
| Goa | 15.755 | 72.819 | 503 | 0.04355 |
| Goa | 15.707 | 73.009 | 205 | 0.07799 |
| Goa | 15.616 | 73.228 | 93 | 1.64047 |
| Kochi | 10.176 | 75.907 | 94 | 0.35533 |
| Kochi | 10.078 | 75.634 | 250 | 0.35533 |
| Kochi | 10.044 | 75.529 | 500 | 0.35533 |
| Mangalore | 12.826 | 74.339 | 102 | 0.6854 |
| Mangalore | 13.058 | 74.071 | 176 | 0.14161 |
| Mangalore | 13.005 | 74.012 | 983 | 0.47906 |
| Mangalore | 13.005 | 74.012 | 200 | 0.13913 |
| Trivandrum | 8.649 | 76.662 | 110 | 0.38569 |
| Trivandrum | 8.724 | 76.428 | 196 | 0.34403 |
| Trivandrum | 8.626 | 76.229 | 500 | 0.46625 |
| Trivandrum | 8.626 | 76.229 | 750 | 0.46665 |

**Table I** Details of sampling site and Physico-chemical parameters of samples collected from A) 492 Arabian Sea and B) Bay of Bengal waters.
| Location | Latitude | Longitude | Depth (m) | Oxygen concentration (ml/L) |
| --- | --- | --- | --- | --- |
| Station1 | 1956.762N | 8652.630E | 70 | 0.55 |
| Station2 | 1956.762N | 8652.630E | 70 | 0.55 |
| Station3 | 1956.762N | 8652.630E | 70 | 0.55 |
| Station4 | 1956.762N | 8652.630E | 70 | 0.55 |
| Station5 | 1921.658N | 8546.100E | 510 | 0.07 |
| Station6 | 1707.369N | 8328.174E | 40 | 0.53 |
| Station7 | 1707.369N | 8328.174E | 40 | 0.53 |
| Station8 | 1956.762N | 8652.630E | 320 | 0.03 |
| Station9 | 1508.014N | 8033.868E | 912 | 0.52 |
| Station10 | 1508.014N | 8033.868E | 912 | 0.52 |
| Station11 | 1811.692N | 8427.742E | 468 | 0.04 |
| Station12 | 1707.369N | 8328.174E | 302 | 0.05 |
| Station13 | 1707.369N | 8328.174E | 302 | 0.05 |
| Station14 | 1557.937N | 8142.164E | 359 | 0.11 |
| Station15 | 1800.468N | 8421.505E | 408 | 0.07 |
| Station16 | 1800.468N | 8421.505E | 408 | 0.07 |
| Station17 | 1931.112N | 8553.380E | 550 | 0.05 |
| Station18 | 1931.112N | 8553.380E | 550 | 0.05 |
| Station19 | 1508.014N | 8033.868E | 633 | 0.09 |
| Station20 | 1508.014N | 8033.868E | 633 | 0.09 |
| Station21 | 1557.937N | 8142.164E | 910 | 0.41 |
| Station22 | 1947.570N | 8649.754E | 509 | 0.05 |
| Station23 | 1947.570N | 8649.754E | 509 | 0.05 |
| Station24 | 1707.369N | 8328.174E | 302 | 0.05 |
| Station25 | 1707.369N | 8328.174E | 302 | 0.05 |
| Station26 | 1947.570N | 8649.754E | 20 | 0.623 |
| Station27 | 1947.570N | 8649.754E | 20 | 0.623 |
| Station28 | 1800.468N | 8421.505E | 817 | 0.28 |
| Station29 | 1800.468N | 8421.505E | 817 | 0.28 |
| Station30 | 1921.658N | 8546.100E | 340 | 0.074 |
| Station31 | 1921.658N | 8546.100E | 340 | 0.074 |

The bacteria were characterized and subjected to16S rRNA sequencing followed by BLASTn homology search. All these 16S rRNA sequences were deposited in NCBI GenBank under accession numbers MH217583, MH256039 to MH256108 (Table II). Each 16S rRNA was subjected to MEGA6 analysis with 10 iterations and 1000 bootstraps to identify phylogenetic relationship. 16S rRNA based analysis has identified 71 bacteria belonging to genus *Halomonas*, *Marinobacter*, *Idiomarina*, *Pshyctobacter*, *Pseudoalteromonas* along with two more which are represented only by the single sequences those were *Thallasospira* (in Bay of Bengal) and *Paenibacillus* (in Arabian sea). In case of both Arabian Sea and Bay of Bengal the distribution was not specific with respect to either depth, collection site and oxygen concentrations. For example, bacteria from genus *Halomonas* or *Pseudoalteromonas* could be isolated from almost all samples in BOB and AS waters. However, some bacteria like *Thallasospira* was observed only at BOB waters and bacteria from genus *Paenibacillus* was isolated only from AS. Thus, the initial 16SrRNA analysis revealed that from AS in total 6 genera while from BOB bacteria belonging to 2 genera were successfully cultured, indicating greater diversity in AS than BOB.

**Table II.**
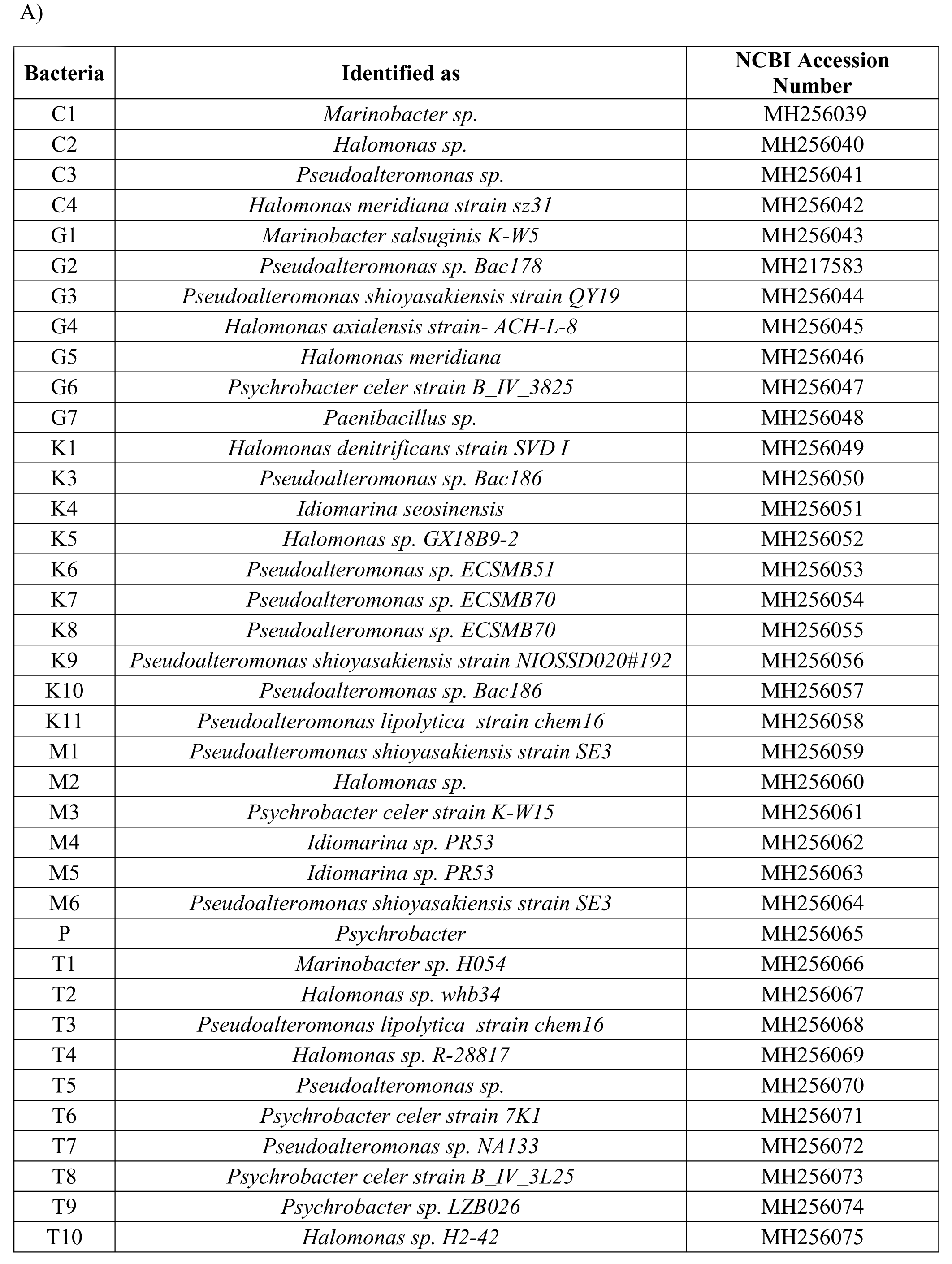
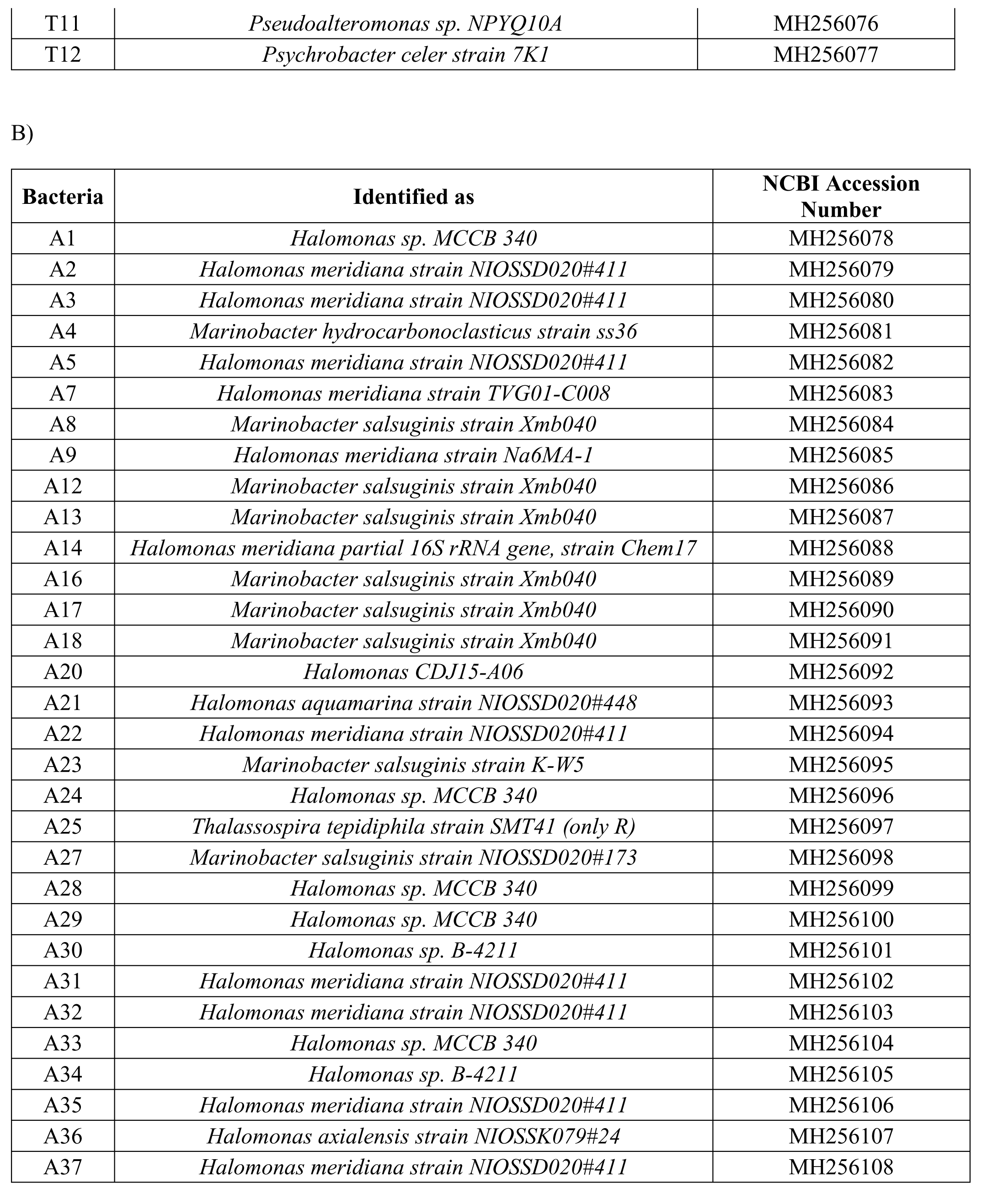
Identified bacteria and corresponding NCBI gene sequence ID for A) Arabian sea B) Bay of Bengal

| **Bacteria** | **Identified as** | **NCBI Accession Number** |
| --- | --- | --- |
| C1 | *Marinobacter sp.* | MH256039 |
| C2 | *Halomonas sp.* | MH256040 |
| C3 | *Pseudoalteromonas sp.* | MH256041 |
| C4 | *Halomonas meridiana strain sz31* | MH256042 |
| G1 | *Marinobacter salsuginis K-W5* | MH256043 |
| G2 | *Pseudoalteromonas sp. Bac178* | MH217583 |
| G3 | *Pseudoalteromonas shioyasakiensis strain QY19* | MH256044 |
| G4 | *Halomonas axialensis strain- ACH-L-8* | MH256045 |
| G5 | *Halomonas meridiana* | MH256046 |
| G6 | *Psychrobacter celer strain B_IV_3825* | MH256047 |
| G7 | *Paenibacillus sp.* | MH256048 |
| K1 | *Halomonas denitrificans strain SVD I* | MH256049 |
| K3 | *Pseudoalteromonas sp. Bac186* | MH256050 |
| K4 | *Idiomarina seosinensis* | MH256051 |
| K5 | *Halomonas sp. GX18B9-2* | MH256052 |
| K6 | *Pseudoalteromonas sp. ECSMB51* | MH256053 |
| K7 | *Pseudoalteromonas sp. ECSMB70* | MH256054 |
| K8 | *Pseudoalteromonas sp. ECSMB70* | MH256055 |
| K9 | *Pseudoalteromonas shioyasakiensis strain NIOSSD020#192* | MH256056 |
| K10 | *Pseudoalteromonas sp. Bac186* | MH256057 |
| K11 | *Pseudoalteromonas lipolytica strain chem16* | MH256058 |
| M1 | *Pseudoalteromonas shioyasakiensis strain SE3* | MH256059 |
| M2 | *Halomonas sp.* | MH256060 |
| M3 | *Psychrobacter celer strain K-W15* | MH256061 |
| M4 | *Idiomarina sp. PR53* | MH256062 |
| M5 | *Idiomarina sp. PR53* | MH256063 |
| M6 | *Pseudoalteromonas shioyasakiensis strain SE3* | MH256064 |
| P | *Psychrobacter* | MH256065 |
| T1 | *Marinobacter sp. H054* | MH256066 |
| T2 | *Halomonas sp. whb34* | MH256067 |
| T3 | *Pseudoalteromonas lipolytica strain chem16* | MH256068 |
| T4 | *Halomonas sp. R-28817* | MH256069 |
| T5 | *Pseudoalteromonas sp.* | MH256070 |
| T6 | *Psychrobacter celer strain 7K1* | MH256071 |
| T7 | *Pseudoalteromonas sp. NA133* | MH256072 |
| T8 | *Psychrobacter celer strain B_IV_3L25* | MH256073 |
| T9 | *Psychrobacter sp. LZB026* | MH256074 |
| T10 | *Halomonas sp. H2-42* | MH256075 |
| T11 | *Pseudoalteromonas sp. NPYQ10A* | MH256076 |
| T12 | *Psychrobacter celer strain 7K1* | MH256077 |

**Table II** Identified bacteria and corresponding NCBI gene sequence ID for A) Arabian sea B) Bay 500 of Bengal
| **Bacteria** | **Identified as** | **NCBI Accession Number** |
| --- | --- | --- |
| A1 | *Halomonas sp. MCCB 340* | MH256078 |
| A2 | *Halomonas meridiana strain NIOSSD020#411* | MH256079 |
| A3 | *Halomonas meridiana strain NIOSSD020#411* | MH256080 |
| A4 | *Marinobacter hydrocarbonoclasticus strain ss36* | MH256081 |
| A5 | *Halomonas meridiana strain NIOSSD020#411* | MH256082 |
| A7 | *Halomonas meridiana strain TVG01-C008* | MH256083 |
| A8 | *Marinobacter salsuginis strain Xmb040* | MH256084 |
| A9 | *Halomonas meridiana strain Na6MA-1* | MH256085 |
| A12 | *Marinobacter salsuginis strain Xmb040* | MH256086 |
| A13 | *Marinobacter salsuginis strain Xmb040* | MH256087 |
| A14 | *Halomonas meridiana partial 16S rRNA gene, strain Chem17* | MH256088 |
| A16 | *Marinobacter salsuginis strain Xmb040* | MH256089 |
| A17 | *Marinobacter salsuginis strain Xmb040* | MH256090 |
| A18 | *Marinobacter salsuginis strain Xmb040* | MH256091 |
| A20 | *Halomonas CDJ15-A06* | MH256092 |
| A21 | *Halomonas aquamarina strain NIOSSD020#448* | MH256093 |
| A22 | *Halomonas meridiana strain NIOSSD020#411* | MH256094 |
| A23 | *Marinobacter salsuginis strain K-W5* | MH256095 |
| A24 | *Halomonas sp. MCCB 340* | MH256096 |
| A25 | *Thalassospira tepidiphila strain SMT41 (only R)* | MH256097 |
| A27 | *Marinobacter salsuginis strain NIOSSD020#173* | MH256098 |
| A28 | *Halomonas sp. MCCB 340* | MH256099 |
| A29 | *Halomonas sp. MCCB 340* | MH256100 |
| A30 | *Halomonas sp. B-4211* | MH256101 |
| A31 | *Halomonas meridiana strain NIOSSD020#411* | MH256102 |
| A32 | *Halomonas meridiana strain NIOSSD020#411* | MH256103 |
| A33 | *Halomonas sp. MCCB 340* | MH256104 |
| A34 | *Halomonas sp. B-4211* | MH256105 |
| A35 | *Halomonas meridiana strain NIOSSD020#411* | MH256106 |
| A36 | *Halomonas axialensis strain NIOSSK079#24* | MH256107 |
| A37 | *Halomonas meridiana strain NIOSSD020#411* | MH256108 |

### 3.2. Characteristics of bacteria

*Halomonas spp.* has been isolated from hydrocarbon contaminated industrial brine [40], sea water [41], hypersaline lake, hydrothermal vent, saltern fermented food. Members of this genus are involved in bioremediation process [41], some *spp*. can produce biosurfactants. We could identify different *Halomonas* species such as *Halomonas aquamarina, Halomonas axialensis, Halomonas meridiana* and *Halomonas denitrificans* from different location and water depths. Few of these bacteria were reported to have specific characteristics that can be useful in various applications. Such as *Halomonas axialensis* has been shown to reduce nitrate in both aerobic and anaerobic condition and is also oxidase positive [42]. It is known to harbor 20 aldehyde dehydrogenase genes involved in 14 metabolic pathways of aldehyde degradation[43]. *Halomonas meridiana* has genes such as choline dehydrogenase and betaine aldehyde dehydrogenase which are responsible for production of osmo-protectants. Genes responsible for osmolyte, ecotine production (ectABC) and for utilization of ecotine (eutED) have also reported in its genome [44].

Many bacteria from *Marinobacter* genus were also isolated from BOB and AS waters. Interestingly, many species from *Marinobacter* genus are known to degrade petroleum and a few can tolerate metalloids. One of the isolate was identified as *Marinobacter hydrocarbonoclasticus* which is known to grow on aromatic hydrocarbons and can produce extracellular surface-active compound by forming biofilm on n-alkanes, fatty alcohols, apolar lipids, wax esters and triglycerides. Another isolate *Thalassospira tepidiphila* was initially found in petroleum-contaminated seawater and was reported to degrade polycyclic aromatic hydrocarbons. It also shows degradation of naphthalene, phenanthrene, dibenzothiophene and fluorene (mixed into crude oil) [45]. We have also isolated *Pseudoalteromonas shioyasakiensis* which was reported to show halo-alkali and thermo tolerant propertied in addition to chitinase production [46]. Same spp. isolated from Indonesian marine areas, shows polyaromatic hydrocarbon degradation activity. In current report, some of the isolates of *Pseudoalteromonas* were seen to possess hydrocarbon degradation property. Few microbes from *Idiomarina* genus were also identified and bacteria from this genus are known for their chitin degradation activity and production of biodiesel from Jatropha oil [47]. Thus, microbes isolated from Arabian Sea and Bay of Bengal belong to various genus and species which were reported to have many industrial and biological applications. Therefore these isolates were further assessed for various chemical and enzymatic properties.

### 3.3. Carbohydrate utilizing ability of bacteria

Carbohydrates are important part of a global carbon cycle. Marine surface water contains up to 21% carbohydrates in the form of dissolved organic carbon. Polysaccharides and monosaccharide are reported to be present at for 3-4 μM concentration in Pacific, Atlantic and Antarctic Ocean [48]. Carbohydrate utilization is a central pathway for energy production in bacteria (Aluwihare and Repeta, 1999; Arnosti, 2000, Zoppini et al, 2010). Marine heterotrophic bacteria contribute carbon cycle either by the remineralisation of organic carbon or by the production of new bacterial biomass.

Previous reports have demonstrated the presence of carbohydrate degrader’s, belonging to *Bacillus*, *Vibrio, Marinobacter,* etc from estuarine sources near Arabian sea [45]. The characteristic ability of microorganisms to utilise different carbon sources is important in the understanding the marine microbial ecology [49]. Among the bacterial isolates, most of the bacteria were able to utilize multiple carbohydrates while only a few bacteria (C1 from AS and A13, A16 and A23 from BOB) were unable utilize any carbohydrate (Online resource 3). Further, amongst Bay of Bengal isolates, citrate was preferentially used followed by malonate (Fig 1). While Arabian Sea isolates were observed to utilize more complex carbohydrates such as esculin and malonate (Online resource 3). This underlines the basic difference between energy production pathways and complexity of the bacterial metabolic abilities between various isolates from two different Indian oceans. Further a few microbes were observed to utilize only one carbon source, namely, A1 (*Halomonas sp. MCCB 340*), A3 (*Halomonas meridiana* strain NIOSSD020#411), A4 (*Marinobacter hydrocarbonoclasticus* strain ss36) from BOB utilized citrate as a carbon source (Online resource 3) while G3 (*Pseudoalteromonas shioyasakiensis* strain QY19), K10 (*Pseudoalteromonas* sp. Bac186), M1 (*Pseudoalteromonas shioyasakiensis* strain SE3), M2 (*Halomonas* sp.), M4 (*Idiomarina* sp. PR53), M6 (*Pseudoalteromonas shioyasakiensis* strain SE3) and T2 *(Halomonas sp. whb34*) could use single carbohydrate for energy production (Online resource 3). Thus, it could be observed that similar type of bacteria isolated from two different location or conditions have entirely different preferences for carbon sources and energy production pathways.

### 3.4. Enzyme assays

Ocean contains several carbon sources as nutrients. Most of the primary production in marine environments goes into the detrital food which is consumed by heterotrophic microorganisms. The complex macromolecular detritus must be degraded initially into simpler substrates by enzymes which can be utilized by other living organisms. These extracellular enzymes are released into the environment either by secretion or to some extent by cell lysis [50, 51]. In addition to microbial diversity, the types of extracellular enzymes, their magnitude and nature can reflect the probable consumption and diversity of available organic matter in a particular ecosystem. Both aerobic and anaerobic bacteria are known to produce such extracellular enzymes degrade organic matter [52]. Aerobic heterotrophic bacteria often possess degradative pathways for structurally complex organic matter [52–54] and thus can play a significant role in maintenance of nutrient cycling. From this study, we report that in both Bay of Bengal and Arabian Sea amylase, agarase producers and hydrocarbon utilizing micro-organisms are predominant in number. Among the 71 isolates, 42 isolates showed amylase activity (Fig 2 Online resource 4) and 17 isolates showed agarase activity (Fig 2, Online resource 4).

Further, 6 isolates from BOB (Fig 2, Online resource 4) and 28 from AS (Fig 2, Online resource 4) also showed cellulose utilizing capability (Fig 2). In comparison, microbes with chitinase and xylanse activity were less in number (Fig 2, Online resource 4). Groups of the micro-organisms known as the hydrocarbonoclastic bacteria (HCB) play an important role in the biodegradation and removal of petrochemical pollutants in the global ocean. Oil spill is one of the serious issues among the contaminants in the ocean where the presence and activities of these bacteria has been detected. Oil (hydrocarbons) is spilled in the ocean from the ships. Thus, in addition to bacterial diversity, evaluating the ability of bacteria to utilize various organic substrates is an important aspect to understand the functional marine ecology. Over 175 genera, distributed across several major bacterial classes such as Alpha, Beta, and Gammaproteobacteria, Actinomycetes, Flavobacteria include representatives of HCB species [55]. Out of 71 isolates, 32 bacteria showed promising hydrocarbon utilization activity (Fig 2, Online resource 4).

In OMZ regions, organisms are known to use nitrate and sulfate as a terminal electron acceptor and as one of the energy sources. 3 isolates from AS demonstrated sulfur reduction ability (Fig 2, Online resource 4) whereas 2 from BOB and 3 from AS showed nitrate reduction activity (Fig 2, Online resource 4). Among all the isolates, very few isolates show multiple enzymatic activities. A1 (*Halomonas sp*. MCCB 340) from Bay of Bengal was hydrocarbon degrading as well as had nitrate and sulfur reducing activity (Fig 2, Online resource 4). Enzymatic activities of each isolates from BOB & AS are summarized (Fig 2, Online resource 4). A recent study from the Arabian sea, also suggested that some bacteria are capable for complete nitrate reduction [56]. Bacteria of from genera such as *Halomonas* and *Pseudoalteromonas* are known to be involved in nitrogen cycle [57, 58]. *Pseudoalteromonas*, *Halomonas* and *Paenibacillus* speices are well known for nitrate reduction, with the presences of NirS gene [59–61]. In addition, presences of denitrification process, supporting the presences of denitrifier microbial populations in BOB and Arabian sea regions, is recently reported [28, 56]. Also, a metagenomics analysis of OMZ and non-OMZ regions of Bay Bengal suggests the difference in bacterial abundance. Functional analysis revealed the presence of assimilatory Sulphur reducing genes in the non-OMZ areas whereas the dissimilatory sulphate reducing genes were abundant in OMZ regions. Also, our study has revealed the distinct bacterial diversity of Bay of Bengal when compared to OMZ regions of Peru and Chile. This work also reports the presence of bacteria involved in nitrogen metabolism [62]. Therefore, current analysis with support from the previous studies confirms the presences of nitrate reducing cultivable bacteria in BOB and Arabian Sea areas.

### 3.5. Antibiotic assay

Bacteria acquire antibiotic resistance naturally or through horizontal gene transfer from other organisms and by contact with antimicrobial agents. Many of the isolates were found to resist higher concentration (100μg/mL) of ampicillin, chloramphenicol, kanamycin and streptomycin antibiotics. Ampicillin susceptibility to 100μg/mL concentration was only shown by 9 AS bacteria and 20 BOB bacteria (Fig 3, Online resource 5) and a few were weakly resist 100μg/mL concentration of ampicillin and chloramphenicol respectively (Fig 3, Online resource 5). Susceptibility to 100μg/mL concentration of chloramphenicol was shown by 11 bacterial isolates from AS and 1 bacterial isolate (BOB). Resistance to 100μg/mL kanamycin and streptomycin was detected in all 71 isolates (Fig 3). Therefore, our results revealed that AS and BOB is reservoir of chloramphenicol, kanamycin and streptomycin resistant bacteria and this genetic trait is predominant in these isolates.

The OMZ and Non-OMZ areas of the Indian Ocean thus harbor diverse bacteria with different catabolic efficiencies. In conclusion, current study has identified 71 distinct bacteria from AS and BOB water samples which differ in dissolved oxygen concentration. The identified bacteria belong to different genera and species which have varied physical and biochemical properties including carbohydrate utilization, hydrocarbon utilization. The supply of organic material to the OMZ and Non-OMZ water column is an important factor as macro- as well as micro-life forms survive on this organic matter. Therefore, future studies including detailed investigation of currently identified cultivable bacteria and their potential applications and exploration of more bacterial diversity from Indian Ocean is warranted and is likely to provide rich and diverse microbial populations. Also, this study identifies the denitrifying microbial population which will be useful in studying the nitrogen cycle in the Arabian Sea and Bay of Bengal. Overall, the current study provides basic identification of bacterial diversity and their characterization in Indian oceans.

## Acknowledgments

This work was funded by Ministry of Earth Sciences (MoES), Government of India, under the Microbial Oceanography scheme. The funding agency had no role in study design, data collection and analysis, decision to publish, or preparation of the manuscript. The authors acknowledge support from Dr. Somasundar of MOES.

## Author contributions

DDD, SNR and MP designed the experiments; YMP, RB, AMK and PGM carried out the wet lab work; SNR, YMP, RB, carried out bioinformatics and statistical analysis, DDD and SNR interpreted the results and wrote the manuscript.

## Funding and Acknowledgments

This work was funded by Ministry of Earth Sciences (MoEs), Government of India, under the Microbial Oceanography scheme to DDD. The funding agency had no role in study design, data collection and analysis, decision to publish, or preparation of the manuscript. Authors acknowledge Mr. Suraj Joshi, who helped in water sample collection. This project was coordinated through CMLRE and we acknowledge the help and support from Dr. Saravannane, Dr. Cherayi, Dr. Shivaji and Director of CMLRE, Dr. Sudhakar.

## Figure Legends

**Figure 1** Several carbohydrate utilizing capacities of bacterial isolates from Arabian sea and Bay of Bengal.

**Figure 2** Bacterial isolates capable to produce various substrate degrading enzymes.

**Figure 3** Number of bacterial isolates showing resistances to different antibiotics.

## Supplementary Material Legends

**Supplementary material 1** List of primers used in this study

**Supplementary material 2** Colony characterization of AS and BoB isolates

**Supplementary material 3** Carbohydrates utilized by AS and BoB isolates

**Supplementary material 4** Enzyme activity shown by AS and BoB isolates

**Supplementary material 5** Antibiotic resistance shown by AS and BoB isolates

## References

1. Satheesh S, Ba-akdah MA, Al-Sofyani AA (2016) Natural antifouling compound production by microbes associated with marine macroorganisms: A review. Electronic Journal of Biotechnology 19:26–35

2. Armstrong E, Yan L, Boyd KG, et al (2001) The symbiotic role of marine microbes on living surfaces. Hydrobiologia 461:37–40

3. Debbab A, Aly AH, Lin WH, Proksch P (2010) Bioactive compounds from marine bacteria and fungi. Microbial biotechnology 3:544–563

4. Thakur AN, Thakur NL, Indap MM, et al (2005) Antiangiogenic, antimicrobial, and cytotoxic potential of sponge-associated bacteria. Marine Biotechnology 7:245–252

5. Zheng L, Han X, Chen H, et al (2005) Marine bacteria associated with marine macroorganisms: the potential antimicrobial resources. Annals of Microbiology 55:119–124

6. Hu Y, Chen J, Hu G, et al (2015) Statistical Research on the Bioactivity of New Marine Natural Products Discovered during the 28 Years from 1985 to 2012. Marine Drugs 13:202–221. https://doi.org/10.3390/md13010202

7. Lam P, Kuypers MMM (2011) Microbial Nitrogen Cycling Processes in Oxygen Minimum Zones. Annual Review of Marine Science 3:317–345. https://doi.org/10.1146/annurev-marine-120709-142814

8. Meyerhof MS, Wilson JM, Dawson MN, Michael Beman J (2016) Microbial community diversity, structure and assembly across oxygen gradients in meromictic marine lakes, Palau: Microbial communities in meromictic marine lakes. Environmental Microbiology 18:4907–4919. https://doi.org/10.1111/1462-2920.13416

9. Anas A, Nilayangod C, Jasmin C, et al (2016) Diversity and bioactive potentials of culturable heterotrophic bacteria from the surficial sediments of the Arabian Sea. 3 Biotech 6:238

10. Bakun A, Field DB, Redondo-Rodriguez ANA, Weeks SJ (2010) Greenhouse gas, upwelling-favorable winds, and the future of coastal ocean upwelling ecosystems. Global Change Biology 16:1213–1228

11. Lavik G, Stührmann T, Brüchert V, et al (2009) Detoxification of sulphidic African shelf waters by blooming chemolithotrophs. Nature 457:581

12. Gibson RN, Atkinson RJA (2003) Oxygen minimum zone benthos: adaptation and community response to hypoxia. Oceanogr Marine Biol Annu Rev 41:1–45

13. Bryant JA, Stewart FJ, Eppley JM, DeLong EF (2012) Microbial community phylogenetic and trait diversity declines with depth in a marine oxygen minimum zone. Ecology 93:1659–1673

14. Divya B, Parvathi A, Bharathi PL, Nair S (2011) 16S rRNA-based bacterial diversity in the organic-rich sediments underlying oxygen-deficient waters of the eastern Arabian Sea. World Journal of Microbiology and Biotechnology 27:2821–2833

15. Divya B, Soumya KV, Nair S (2010) 16SrRNA and enzymatic diversity of culturable bacteria from the sediments of oxygen minimum zone in the Arabian Sea. Antonie van Leeuwenhoek 98:9–18

16. Das S, Lyla PS, Khan SA (2007) Spatial variation of aerobic culturable heterotrophic bacterial population in sediments of the continental slope of western Bay of Bengal. Indian Journal of Marine Sciences 36:51–58

17. De J, Sarkar A, Ramaiah N (2006) Bioremediation of toxic substances by mercury resistant marine bacteria. Ecotoxicology 15:385–389

18. Nithya C, Pandian SK (2010) Isolation of heterotrophic bacteria from Palk Bay sediments showing heavy metal tolerance and antibiotic production. Microbiological Research 165:578–593

19. Ramesh S, Jayaprakashvel M, Mathivanan N (2006) Microbial status in seawater and coastal sediments during pre-and post-tsunami periods in the Bay of Bengal, India. Marine Ecology 27:198–203

20. Ruban P, Gunaseelan C (2011) Antibiotic resistance of bacteria from Krishna Godavari Basin, Bay of Bengal, India. Environmental and Experimental Biology 9:33–136

21. Santhiya G, Lakshumanan C, Selvin J, Asha D (2011) Microbiological analysis of seawater and sediments in urban shorelines: Occurrence of heavy metals resistance bacteria on Chennai beaches, Bay of Bengal. Microchemical Journal 99:197–202

22. Basu S, Deobagkar DD, Matondkar SP, Furtado I (2013) Culturable Bacterial Flora Associated with the Dinoflagellate Green *Noctiluca miliaris* During Active and Declining Bloom Phases in the Northern Arabian Sea. Microb Ecol 65:934–954. https://doi.org/10.1007/s00248-012-0148-1

23. Paulmier A, Ruiz-Pino D (2009) Oxygen minimum zones (OMZs) in the modern ocean. Progress in Oceanography 80:113–128. https://doi.org/10.1016/j.pocean.2008.08.001

24. Pitcher A, Villanueva L, Hopmans EC, et al (2011) Niche segregation of ammonia-oxidizing archaea and anammox bacteria in the Arabian Sea oxygen minimum zone. The ISME journal 5:1896

25. Wyman M, Hodgson S, Bird C (2013) Denitrifying alphaproteobacteria from the Arabian Sea that express nosZ, the gene encoding nitrous oxide reductase, in oxic and suboxic waters. Applied and environmental microbiology 79:2670–2681

26. Böttcher ME, Schale H, Schnetger B, et al (2000) Stable sulfur isotopes indicate net sulfate reduction in near-surface sediments of the deep Arabian Sea. Deep Sea Research Part II: Topical Studies in Oceanography 47:2769–2783. https://doi.org/10.1016/S0967-0645(00)00048-5

27. Lüke C, Speth DR, Kox MA, et al (2016) Metagenomic analysis of nitrogen and methane cycling in the Arabian Sea oxygen minimum zone. PeerJ 4:e1924

28. Bristow LA, Callbeck CM, Larsen M, et al (2017) N2 production rates limited by nitrite availability in the Bay of Bengal oxygen minimum zone. Nature Geoscience 10:24–29. https://doi.org/10.1038/ngeo2847

29. McCreary JP, Yu Z, Hood RR, et al (2013) Dynamics of the Indian-Ocean oxygen minimum zones. Progress in Oceanography 112–113:15–37. https://doi.org/10.1016/j.pocean.2013.03.002

30. Milliman JD, Meade RH (1983) World-Wide Delivery of River Sediment to the Oceans. The Journal of Geology 91:1–21. https://doi.org/10.1086/628741

31. Milliman JD, Syvitski JPM (1992) Geomorphic/Tectonic Control of Sediment Discharge to the Ocean: The Importance of Small Mountainous Rivers. The Journal of Geology 100:525–544. https://doi.org/10.1086/629606

32. Seitzinger SP (1990) Denitrification In Aquatic Sediments. In: Denitrification in Soil and Sediment. Springer, Boston, MA, pp 301–322

33. Devol AH (1991) Direct measurement of nitrogen gas fluxes from continental shelf sediments. Nature 349:319–321. https://doi.org/10.1038/349319a0

34. Morrison JM, Codispoti LA, Smith SL, et al (1999) The oxygen minimum zone in the Arabian Sea during 1995. Deep Sea Research Part II: Topical Studies in Oceanography 46:1903–1931. https://doi.org/10.1016/S0967-0645(99)00048-X

35. Kulkarni R, Deobagkar D, Zinjarde S (2018) Metals in mangrove ecosystems and associated biota: A global perspective. Ecotoxicology and Environmental Safety 153:215–228. https://doi.org/10.1016/j.ecoenv.2018.02.021

36. Pichard SL, Campbell L, Paul JH (1997) Diversity of the ribulose bisphosphate carboxylase/oxygenase form I gene (rbcL) in natural phytoplankton communities. Appl Environ Microbiol 63:3600–3606

37. Rotthauwe JH, Witzel KP, Liesack W (1997) The ammonia monooxygenase structural gene amoA as a functional marker: molecular fine-scale analysis of natural ammonia-oxidizing populations. Appl Environ Microbiol 63:4704–4712

38. Ueda T, Suga Y, Yahiro N, Matsuguchi T (1995) Remarkable N2-fixing bacterial diversity detected in rice roots by molecular evolutionary analysis of nifH gene sequences. J Bacteriol 177:1414–1417. https://doi.org/10.1128/jb.177.5.1414-1417.1995

39. Jüngst Angelika, Zumft Walter G. (2001) Interdependence of respiratory NO reduction and nitrite reduction revealed by mutagenesis of nirQ, a novel gene in the denitrification gene cluster of Pseudomonas stutzeri. FEBS Letters 314:308–314. https://doi.org/10.1016/0014-5793(92)81495-8

40. Carlson RP, Oshota O, Shipman M, et al (2016) Integrated molecular, physiological and in silico characterization of two Halomonas isolates from industrial brine. Extremophiles 20:261–274. https://doi.org/10.1007/s00792-015-0806-6

41. Gasperotti AF, Studdert CA, Revale S, Herrera Seitz MK (2015) Draft Genome Sequence of Halomonas sp. KHS3, a Polyaromatic Hydrocarbon-Chemotactic Strain. Genome Announcements 3:e00020–15. https://doi.org/10.1128/genomeA.00020-15

42. Jiang J, Pan Y, Hu S, et al (2014) Halomonas songnenensis sp. nov., a moderately halophilic bacterium isolated from saline and alkaline soils. International Journal of Systematic and Evolutionary Microbiology 64:1662–1669. https://doi.org/10.1099/ijs.0.056499-0

43. Ye J, Ren C, Shan X, Zeng R (2016) Draft Genome Sequence of Aldehyde-Degrading Strain Halomonas axialensis ACH-L-8. Genome announcements 4:e00287–16

44. Meyer JL, Dillard BA, Rodgers JM, et al (2015) Draft genome sequence of Halomonas meridiana R1t3 isolated from the surface microbiota of the Caribbean Elkhorn coral Acropora palmata. Standards in Genomic Sciences 10:. https://doi.org/10.1186/s40793-015-0069-y

45. Khandeparker R, Verma P, Meena RM, Deobagkar DD (2011) Phylogenetic diversity of carbohydrate degrading culturable bacteria from Mandovi and Zuari estuaries, Goa, west coast of India. Estuarine, Coastal and Shelf Science 95:359–366. https://doi.org/10.1016/j.ecss.2011.09.004

46. Makhdoumi A, Dehghani-Joybari Z, Mashreghi M, et al (2015) A novel halo-alkali-tolerant and thermo-tolerant chitinase from Pseudoalteromonas sp. DC14 isolated from the Caspian Sea. International Journal of Environmental Science and Technology 12:3895–3904. https://doi.org/10.1007/s13762-015-0848-4

47. Li X, Qian P, Wu S-G, Yu H-Y (2014) Characterization of an organic solvent-tolerant lipase from Idiomarina sp. W33 and its application for biodiesel production using Jatropha oil. Extremophiles 18:171–178. https://doi.org/10.1007/s00792-013-0610-0

48. Wiegmann K, Hensler M, Wohlbrand L, et al (2014) Carbohydrate Catabolism in Phaeobacter inhibens DSM 17395, a Member of the Marine Roseobacter Clade. Applied and Environmental Microbiology 80:4725–4737. https://doi.org/10.1128/AEM.00719-14

49. Giorgio PA del, Cole JJ (1998) Bacterial Growth Efficiency in Natural Aquatic Systems. Annual Review of Ecology and Systematics 29:503–541. https://doi.org/10.1146/annurev.ecolsys.29.1.503

50. Arnosti C (2011) Microbial Extracellular Enzymes and the Marine Carbon Cycle. Annual Review of Marine Science 3:401–425. https://doi.org/10.1146/annurev-marine-120709-142731

51. Sinsabaugh RL, Shah JJF (2012) Ecoenzymatic Stoichiometry and Ecological Theory. Annual Review of Ecology, Evolution, and Systematics 43:313–343. https://doi.org/10.1146/annurev-ecolsys-071112-124414

52. Hall Robert O., Meyer Judy L. (1998) The trophic significance of bacteria in a detritus-based stream food web. Ecology 79:1995–2012. https://doi.org/10.1890/0012-9658(1998)079[1995:TTSOBI]2.0.CO;2

53. Hedges JI, Keil RG (1995) Sedimentary organic matter preservation: an assessment and speculative synthesis. Marine Chemistry 49:81–115. https://doi.org/10.1016/0304-4203(95)00008-F

54. Mayer LM (1994) Relationships between mineral surfaces and organic carbon concentrations in soils and sediments. Chemical Geology 114:347–363. https://doi.org/10.1016/0009-2541(94)90063-9

55. Prince RC, Gramain A, McGenity TJ (2010) Prokaryotic Hydrocarbon Degraders. In: Handbook of Hydrocarbon and Lipid Microbiology. Springer, Berlin, Heidelberg, pp 1669–1692

56. Mulla A, Fernandes G, Menezes L, et al (2017) Diversity of culturable nitrate-reducing bacteria from the Arabian Sea oxygen minimum zone. Deep Sea Research Part II: Topical Studies in Oceanography. https://doi.org/10.1016/j.dsr2.2017.12.014

57. Cai H, Jiao N (2008) Diversity and Abundance of Nitrate Assimilation Genes in the Northern South China Sea. Microb Ecol 56:751–764. https://doi.org/10.1007/s00248-008-9394-7

58. Jones EBG, Sakayaroj J, Suetrong S, et al (2009) Classification of marine Ascomycota, anamorphic taxa and Basidiomycota. Fungal Diversity 35:187

59. Allen AE, Booth MG, Frischer ME, et al (2001) Diversity and detection of nitrate assimilation genes in marine bacteria. Applied and Environmental Microbiology 67:5343–5348

60. Bourbonnais A, Juniper SK, Butterfield DA, et al (2014) Diversity and abundance of Bacteria and *nirS*-encoding denitrifiers associated with the Juan de Fuca Ridge hydrothermal system. Ann Microbiol 64:1691–1705. https://doi.org/10.1007/s13213-014-0813-3

61. Rütting T, Boeckx P, Müller C, Klemedtsson L (2011) Assessment of the importance of dissimilatory nitrate reduction to ammonium for the terrestrial nitrogen cycle. Biogeosciences 8:1779–1791. https://doi.org/10.5194/bg-8-1779-2011

62. Rajpathak SN, Banerjee R, Mishra PG, et al (2018) An exploration of microbial and associated functional diversity in the OMZ and non-OMZ areas in the Bay of Bengal. Journal of Biosciences

