## Supplementary figures and images for "Isolation, identification and functional characterization of cultivable bacteria from Arabian Sea and Bay of Bengal water samples reveals high diversity"

### Figure 2

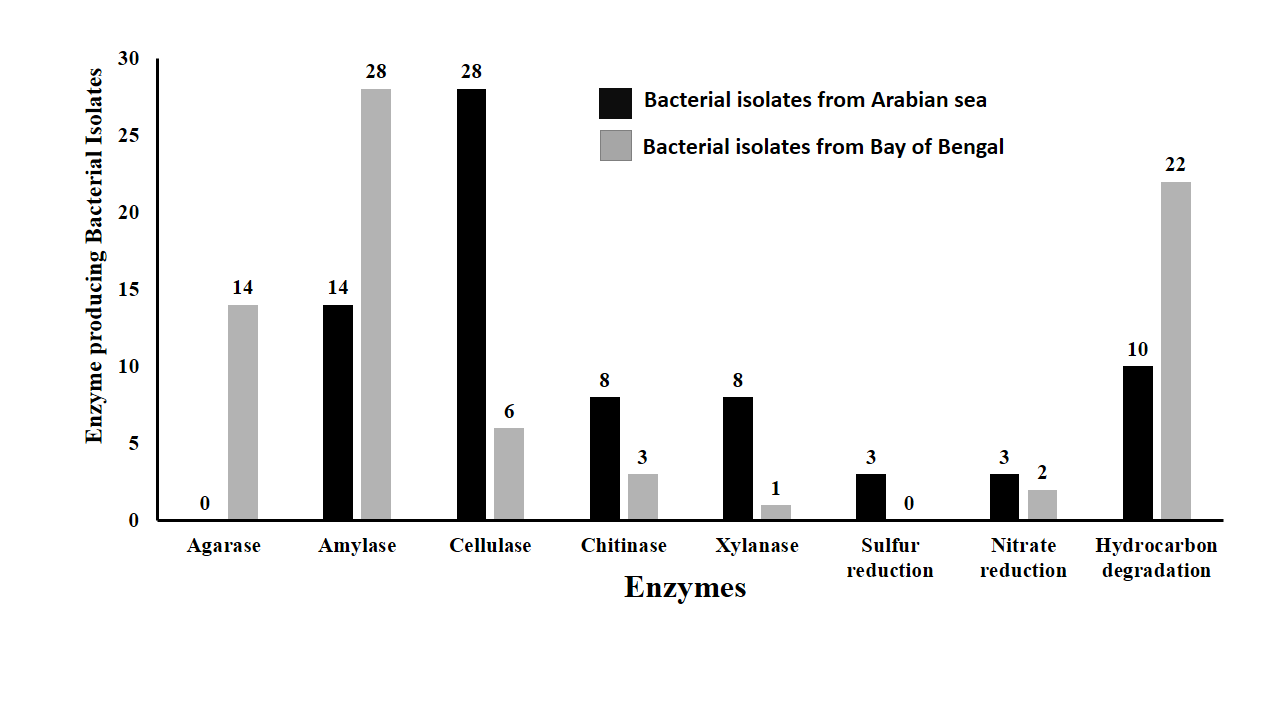

### Figure 3

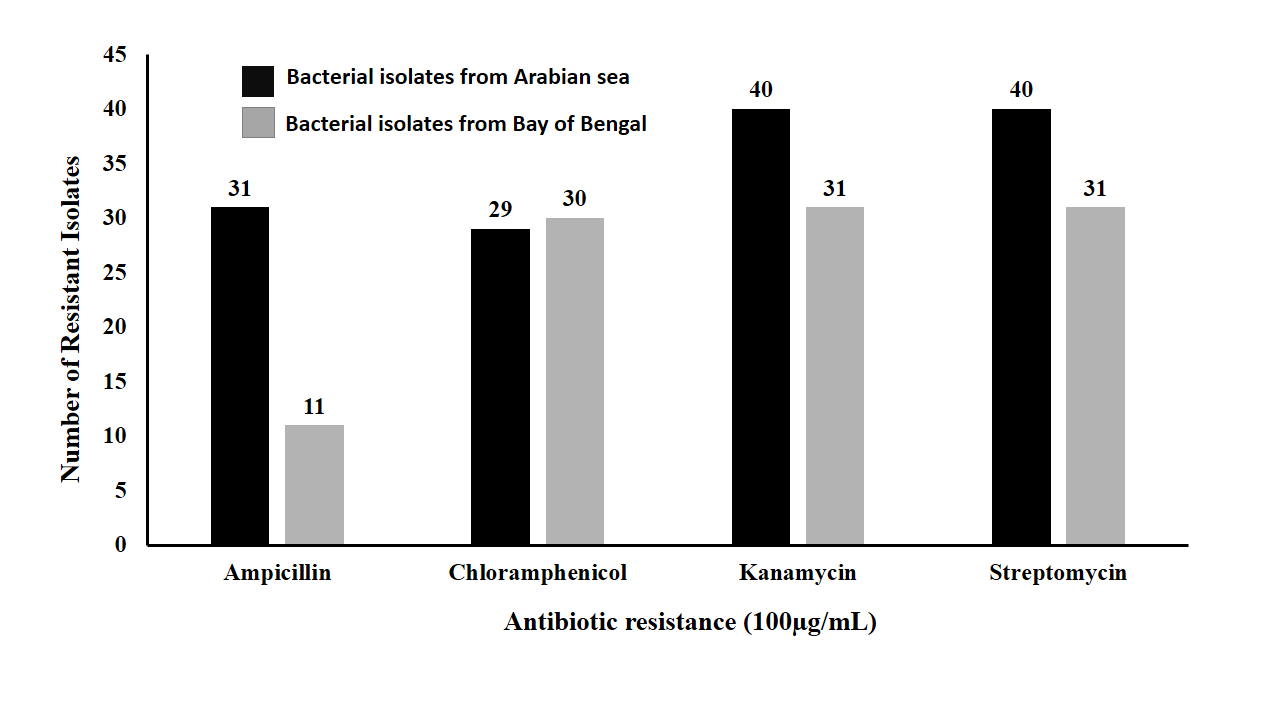
